# The 9aaTAD activation domains in the four Yamanaka Oct4, Sox2, Myc, and Klf4 transcription factors essential during the stem cell development

**DOI:** 10.1101/2019.12.15.876706

**Authors:** Martin Piskacek, Kristina Jendruchova, Martina Rezacova, Marek Havelka, Norbert Gasparik, Alena Hofrova, Andrea Knight

**Affiliations:** Laboratory of Cancer Biology and Genetics, Department of Pathological Physiology, Faculty of Medicine, Masaryk University Brno, Kamenice 5, 625 00 Brno, Czech Republic; Gamma Delta T Cell Laboratory, Department of Pathological Physiology, Faculty of Medicine, Masaryk University Brno, Kamenice 5, 625 00 Brno, Czech Republic

**Keywords:** activation domain, 9aaTAD, phase separation, condensate, Oct4, Pou5f, Sox2, Myc and Klf4

## Abstract

Somatic cells can be reprogrammed by the Yamanaka factors Oct4, Sox2, Myc and Klf4 activators into induced pluripotent stem cells. Throughout their genome, the Oct4, Sox2 and Klf4 cooperate with mediators of transcription, where the DNA binding sites serve as scaffolds for the phase-separated transcriptional condensates at distinct genome loci. In this study, we identified the 9aaTAD activation domains as the common interaction interface of the Yamanaka factors for transcription machinery. All four activation domains were identified by our online 9aaTAD prediction service and experimentally confirmed as strong activators of transcription. We considered the mediator interactions granted by 9aaTADs as part of the Yamanaka factors ability to reprogram cell fate.

**Graphical abstract:** 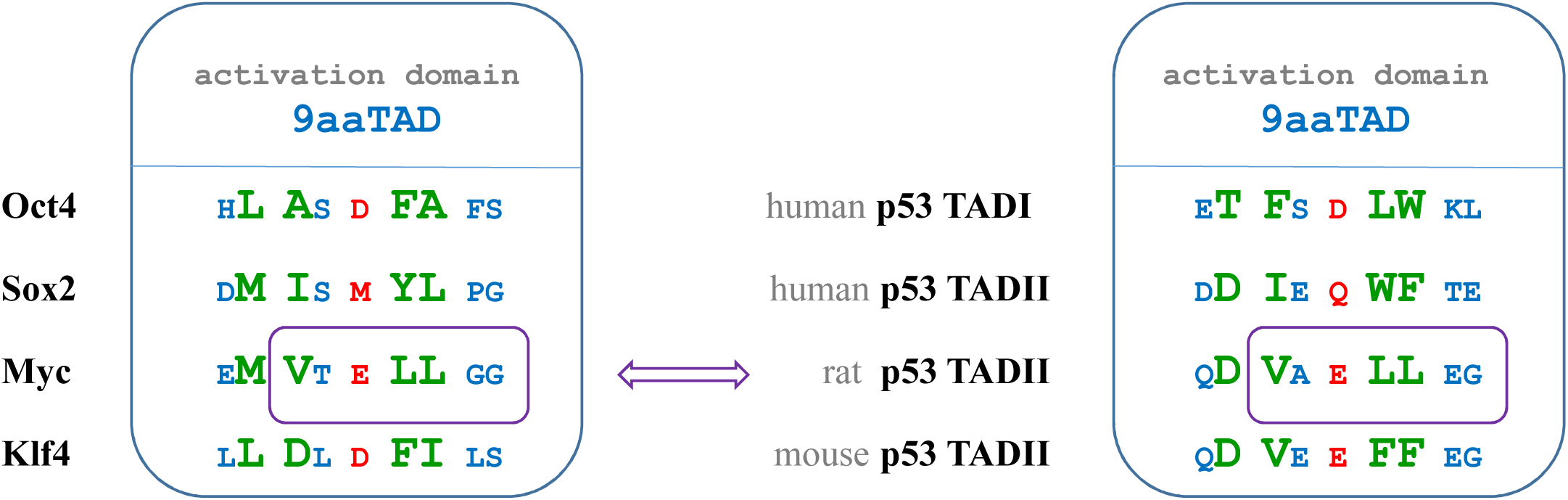

## Introduction

Our longstanding effort has been to determine the Nine-amino-acid TransActivation Domains (9aaTADs) in eukaryotic activators. Previously, we identified the 9aaTAD in a large set of transcription activators that universally recruit multiple mediators of transcription (1–11). The 9aaTAD family is represented by Gal4, p53, MLL, TCF3/E2A and SREBP1. The 9aaTADs were found in the SP/KLF family (Sp1, Sp2, Sp3, Sp4, Klf1, Klf2, Klf3, Klf4, Klf5, Klf6, Klf7, Klf8, Klf12, Klf15, WT1), the SOX family (Sox18 and SoxE), in hormone receptors (RARa, HNF4, PPAR, VDR, NHR49), in yeast transcription factors (Oaf1, Pip2, Pdr1, Pdr3, Rtg3, Gln3, Gcn4, Pho4, Msn2, Msn4, Met4), and in artificial activators of transcription (P201, B42, p53-ECapLL, KBP 2.20, pRJR200, G80BP-A, G80BP-B (12–19).

Although the activation domains have enormous variability, they are universally recognized by transcriptional machinery throughout eukaryotes (20). The 9aaTADs have the competence to activate transcription as small peptides (9 to 14 amino acids long). Currently, all tested human 9aaTADs have been shown to be functional in yeast. Therefore, we considered the 9aaTADs universal function in all eukaryotes as a further ground property of the 9aaTAD family (15, 17, 18, 20, 21). Besides the amino acid pattern, a specific distribution of amino acids in the 9aaTAD is essential. Among the characteristic features is a tandem of hydrophobic clusters surrounded by hydrophilic regions (15, 18). These hydrophobic clusters are often accompanied or are fully substituted by hydrophilic amino acids with aromatic side chains as are tryptophan (W) or tyrosine (Y), e.g. in p53. Accumulation of valines or isoleucines can compromise or fully inactivate the 9aaTAD function (22).

The 9aaTADs are well balanced with hydrophilic amino acids, which may be either positively or negatively charged. Some of them are acidic and negatively charged *e.g.* Gal4 others are positively charged *e.g.* Sp1. The general acidic character of activation domains is old and sadly fixed error, which still persists for Gal4 and also other activators (23). The Gal4 activation domain (DDVY N YLFD) is conserved only in the 9aaTAD region especially in ancestral orthologs as are *Hansenula fabianii* or *Wickerhamomyces ciferrii*, whose activation domain is hardly acidic (NDFY S LIFN)(24). Moreover, we recently demonstrated that the exchange of all negatively charged residues for positively charged ones and *vice versa* in Gal4 and respectively in Sp1 activation domains did not interfere with their function as strong activators of transcription (22).

The 9aaTADs bind to one or more of the mediator’s domains on MED15 or CBP/p300 (1, 23). From structural data for the E2A and MLL activation domains in complex with the KIX domain of CBP mediator, the 9aaTADs form short helices whose lengths vary from 9 to 12 aa (14). Online 9aaTAD prediction (using a residue position matrix search and amino acid clustering) is available on www.piskacek.org. The curated 9aaTADs have been annotated on the protein database UniProt, which now accounts for 145 annotations with 39 human activators (https://www.uniprot.org/uniprot/?query=9aatad&sort=score) (Including the current study, the list of 9aaTAD annotations will be further extended on 165 annotations with 52 human activators).

The Yamanaka transcription factors including the Oct4, Sox2, Myc and Klf4 activators, are all essential for chromatin remodelling and gene activation during the cell reprogramming (25). These activators can reprogram somatic cells into induced pluripotent stem cells (iPSCs) (26). In pluripotent cells, Oct4 / Pou5f from POU family associates with Sox2 to maintain pluripotency or with Sox17 to induce primitive endoderm commitment. The direct interaction between Oct4 and Sox2 is DNA dependent and involves the POU helix A1 from Oct4 and the HMG helix A3 from Sox2 (27). Klf4 cooperates with Oct4 and Sox2 (28, 29) to establish embryonic stem cells (ESCs). Oct4 and Sox2 can form a complex and cooperate during cell reprogramming (30, 31). The activation domain and associated SUMO-interacting motif (SIM, amino acid pattern K/RxE) are required for KLF4 protein stability and are essential for cell pluripotency (32–34).

The SOX family are close relatives of transcription factor SRY (SOX for <u>S</u>RY-related HMG b<u>ox</u> and SRY for <u>s</u>ex-determining region <u>Y</u>) (35–37). The SOX transcription factors are divided into groups A to H and are involved in cell fate determination, development and cancer (38–41).

The most prominent member of the SOX family is Sox2, which supports cellular reprogramming and stem cell pluripotency (40). The Sox2 transactivation is associated with the general TFIID activator complex (42) and with stem cell XPC activator complex (38, 43, 44), with Oct4 (45), with Pax6 (46), with Trim24 (47) and with PARP-1 (48). Sox2 post-translational modifications have been reported to modulate transactivation activity (49–51). Sox2-mediated cell reprogramming could be further enhanced by Sox2 fusion with a strong viral activator VP16 (52). Sox1, Sox3, and Sox15 can replace the function of Sox2 in mouse ES (53, 54).

The gene expression programs, which are controlled by master transcription factors, define the identity of each cell. Recent studies have revealed that master activators form phase-separated condensates with the mediators of transcription at specific genomic loci containing their binding sites (55). The phase separation is a universal cooperative mechanism for transcription, focusing the transcriptional machinery onto DNA enhancer sites, including Oct4, Sox2 and Klf4 binding sites (56, 57). Moreover, activation domains drive nucleosome eviction after activators binding to specific genomic loci (58) and after Oct4 binding, the chromatin accessibility is facilitated by chromatin remodelling factors (59, 60).

In our study, we have focused on the of transcription factors Oct4, Sox2, Myc and Klf4 to determinate their activation interface for the interaction with mediators of transcription, which is a part of cell reprogramming.

## Results

### POU family including Oct4

We have performed our online 9aaTAD prediction for Pou5f / Oct4 (9aaTAD prediction service online, www.piskacek.org) (18) and revealed a single perfect hit to amino acid sequence GHLASDFAF (**Figure 1**). By sequence alignment, we have identified the activation domains in other members of the POU family (**Figure 1**).

**Figure 1.**
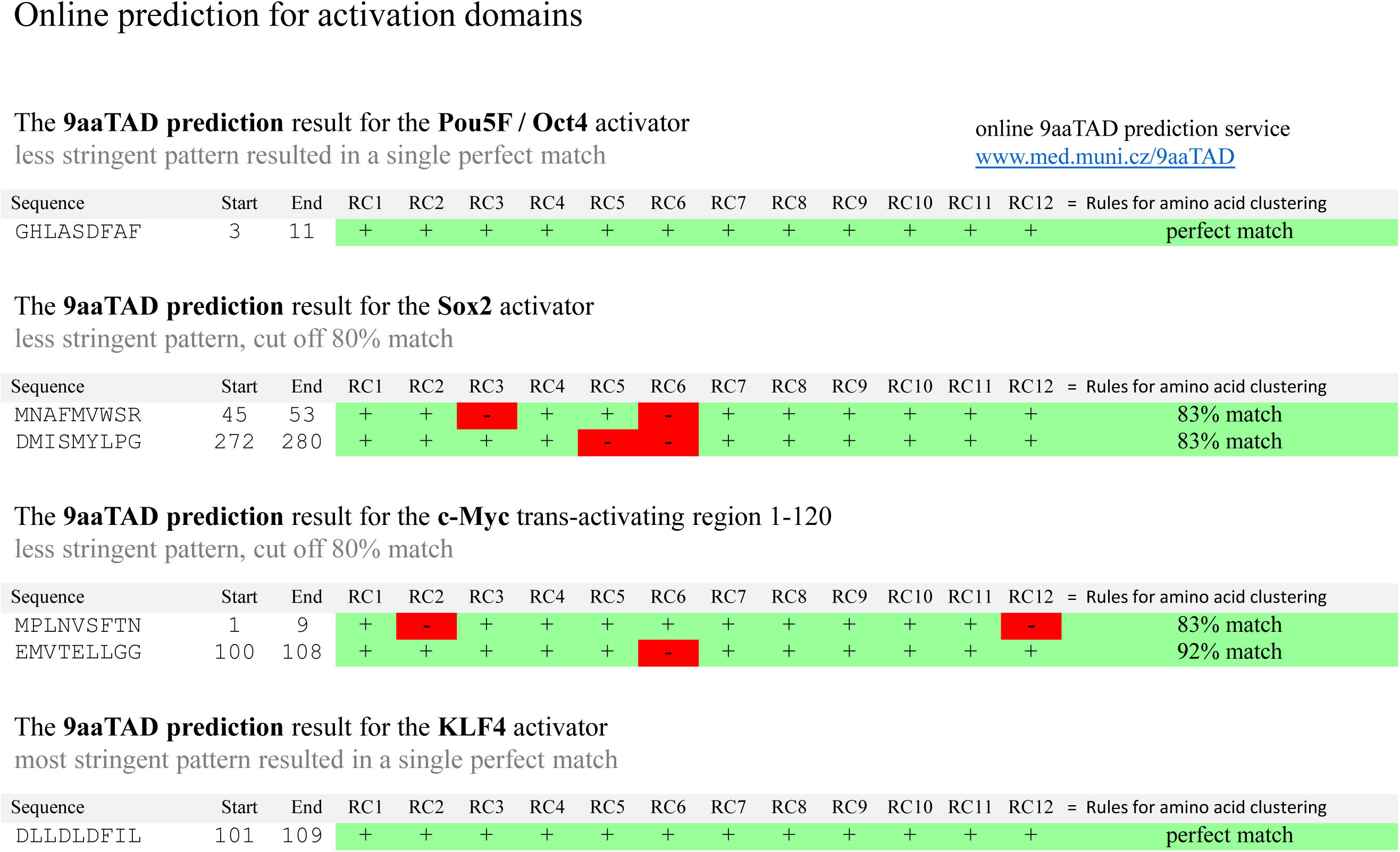
Activation domain prediction. The online 9aaTAD prediction service for activation domain was applied on Pou5F / Oct4, Sox2, Myc (prediction for trans-activation region 50-120 localized experimentally) and Klf4. Algorithm for the 9aaTAD amino acid pattern was applied in the search, and region clustering conformity was assessed by percentage.

We generated LexA constructs, which included the prokaryotic DNA binding domain LexA and POU’s region coding for the 9aaTAD from selected members of clade 1, 3, and 5. The constructs were tested for their ability to activate transcription (**Figure 2**). Both tested human POU activation domains of Pit1 and Pou5f1 have the capacity to activate transcription. Less activity was observed for Pou5 orthologs in Callorhinus ursinus (cur Pou5, Northern fur seal, gene ID: XP_025704146) and Sarcophilus harrisii (shr Pou5, Tasmanian devil, gene ID: G3VJ82), but none in Lingula unguis (lak Pou, brachiopod, gene ID: XP_013392366) or Callorhinchus milii (cmk Pou5, Ghost shark, gene ID: XP_007894073).

**Figure 2.**
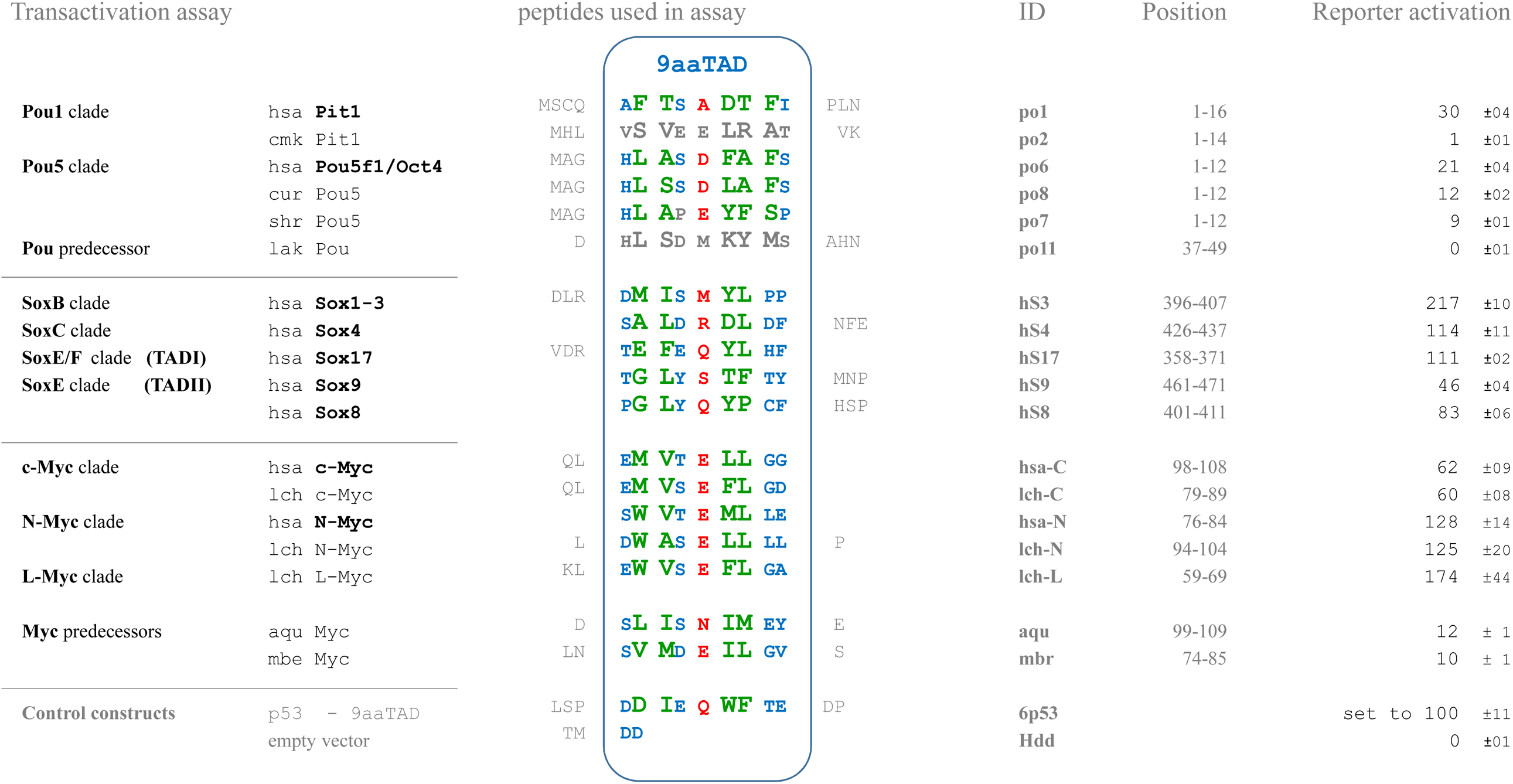
Transactivation Assay. The regions with the identified 9aaTAD activation domains were tested in a reporter assay with hybrid LexA DNA binding domain for the capacity to activate transcription. The average value of the β-galactosidase activities from two independent transformants is presented as a percentage of the reference with standard deviation (means and plusmn; SD; n = 3). We standardized the results to positive control p53 construct 6p53, which was set to 100%. The 9aaTADs activation domains are colored for faster orientation. Deuterated 9aaTADs or their mismatch are in grey. Abbreviations: Homo sapiens (hsa), Callorhinchus milii (cmk, Ghost shark, gene ID: XP_007894073), Callorhinus ursinus (cur Pou5, Northern fur seal, gene ID: XP_025704146), Sarcophilus harrisii (shr Pou5, Tasmanian devil, gene ID: G3VJ82), Lingula unguis (lak Pou, brachiopod, gene ID: XP_013392366), Mnemiopsis leidyi (mle SoxB, comb jelly, gene ID: A0A059XHC3), Amphimedon queenslandica (aqu SoxB, gene ID: B1A9Y6), Strongylocentrotus purpuratus (spu SoxB, purple sea urchin, gene ID: Q9Y0D7 and spu SoxF, gene ID: W4YEI9), Saccoglossus kowalevskii (sko SoxB, hemichordate, common ancestor of chordata, gene ID: Q7YTD4 and sko SoxF, gene ID: B5THP2), *Monosiga brevicollis* (mbe Myc, filozoans, last unicellular ancestor of animals, gene ID: A9V5B4), *Amphimedon queenslandica* (aqu Myc, poriferans, gene ID: XP003390966), *Latimeria chalumnae* (lch c-Myc, lch N-Myc, and lch L-Myc, lobe-finned fish, related to lungfishes and tetrapods, gene IDs: XP005992710, XP005993959, XP006009573).

### SOX family

We have performed online 9aaTAD prediction for Sox2 and revealed two hits above 80% match (**Figure 1**). The first hit was positioned within the DNA binding domain and was therefore excluded. The second hit was located in the Sox2 region 272-280, which corresponds to reported 9aaTAD activation domains in Sox18 (61) and SoxE (12). By sequence alignment, we have identified activation domains in other human members of the SOX family and their distal orthologs from early metazoans, such as Mnemiopsis leidyi (Ctenophora), Amphimedon queenslandica (Porifera), Strongylocentrotus purpuratus (Echinodermata) and Saccoglossus kowalevskii (Hemichordata), which shared significant similarity with Sox2 and other members of the family (**Figure 3**). Importantly, members of the groups B and F have supported the common origin of their activation domains. In contrast, the well-studied paralogs SRY, Sox15 and Sox30 (62–66) have largely diverged and do not share a clear 9aaTAD motif or significant activation domain homology with the aforementioned members of SOX family.

**Figure 3.**
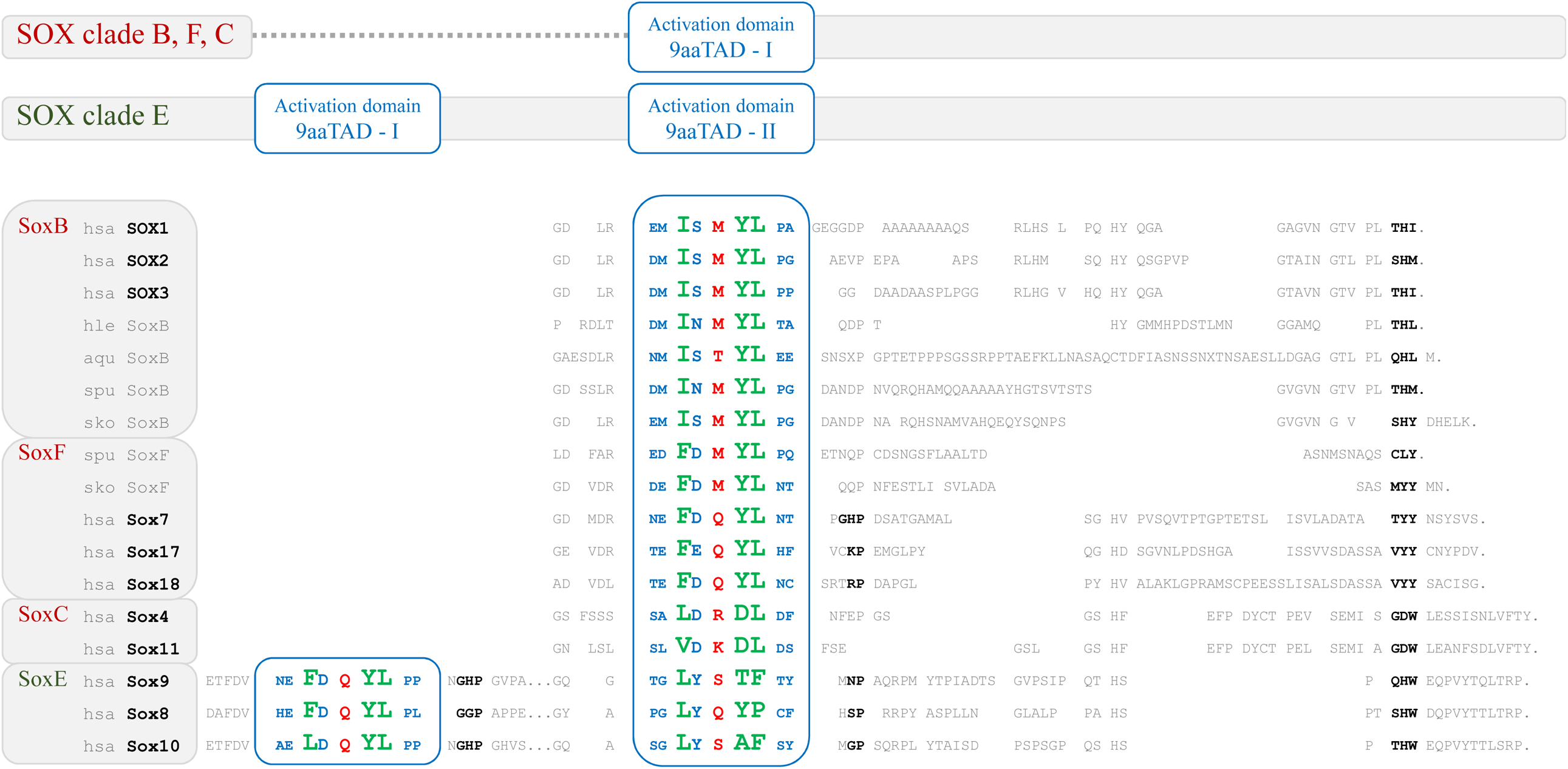
Alignment of SOX family. The C-terminal regions of SOX proteins were aligned by sequence similarities to Sox2 and predicted 9aaTAD activation domain (9aaTAD prediction service online, www.Piskacek.org). The Sox orthologs from early diverged eukaryotes Mnemiopsis leidyi (mle SoxB, gene ID: A0A059XHC3), Amphimedon queenslandica (aqu SoxB, gene ID: B1A9Y6), Strongylocentrotus purpuratus (spu SoxB, gene ID: Q9Y0D7 and spu SoxF, gene ID: W4YEI9) and Saccoglossus kowalevskii (sko SoxB, gene ID: Q7YTD4 and sko SoxF, gene ID: B5THP2) have strong conservation of 9aaTAD. Diversification of the 9aaTAD motif, especially in position p5 (in red) could be seen in human SoxF but not in spu SoxF and sko SoxF. The members of SoxE clade contain two activation domains; the 9aaTADs-I have more conserved sequence but the 9aaTADs-II have more conserved position in the protein and are associated with the SOX C-terminal domains, similarly to the 9aaTADs in all other SOX clades. The 9aaTADs activation domains are colored for faster orientation. Dots in sequences represent stop codon.

We generated LexA constructs, which included the prokaryotic DNA binding domain LexA and SOX’s region coding for the 9aaTAD from selected members of clade B, C, E and F. The constructs were tested for their ability to activate transcription (**Figure 2**). All tested SOX activation domains have the capacity to activate transcription.

### MYC family

We have analysed the 9aaTAD activation domains in MYC activators. The MYC family has multiple conserved regions including MYC boxes MB0 to MBIV and the DNA binding domain (67) (**Figure 4**). From previous studies, the c-Myc region 105-143 was essential for transforming activity (68, 69). Insertion at codon 105 or removal of region 106-143 diminished c-Myc transcriptional activity (70). Based on this, we predicted the activation domain to be localised between MYC boxes MBI and MBII (**Figure 4**).

**Figure 4.**
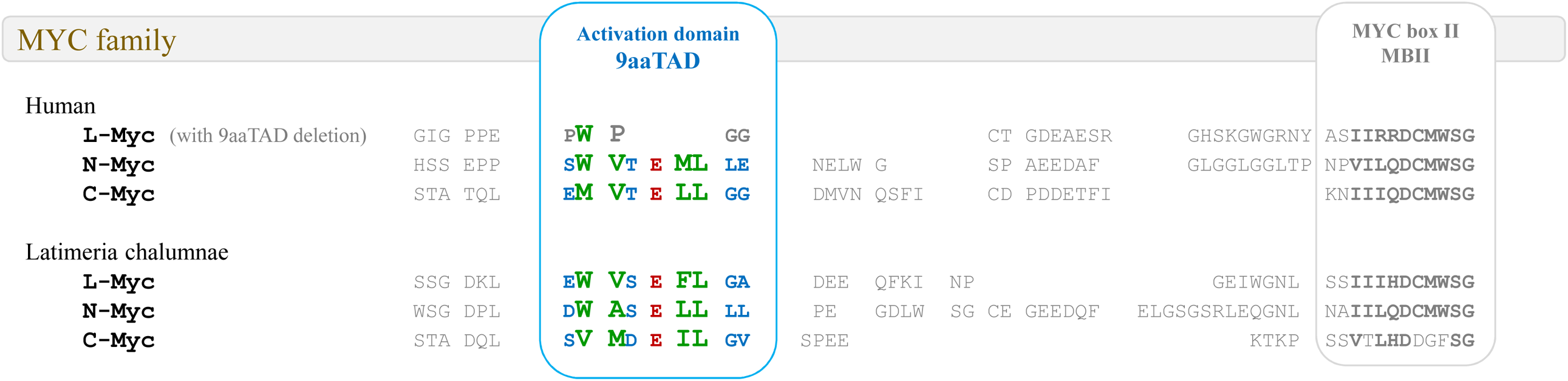
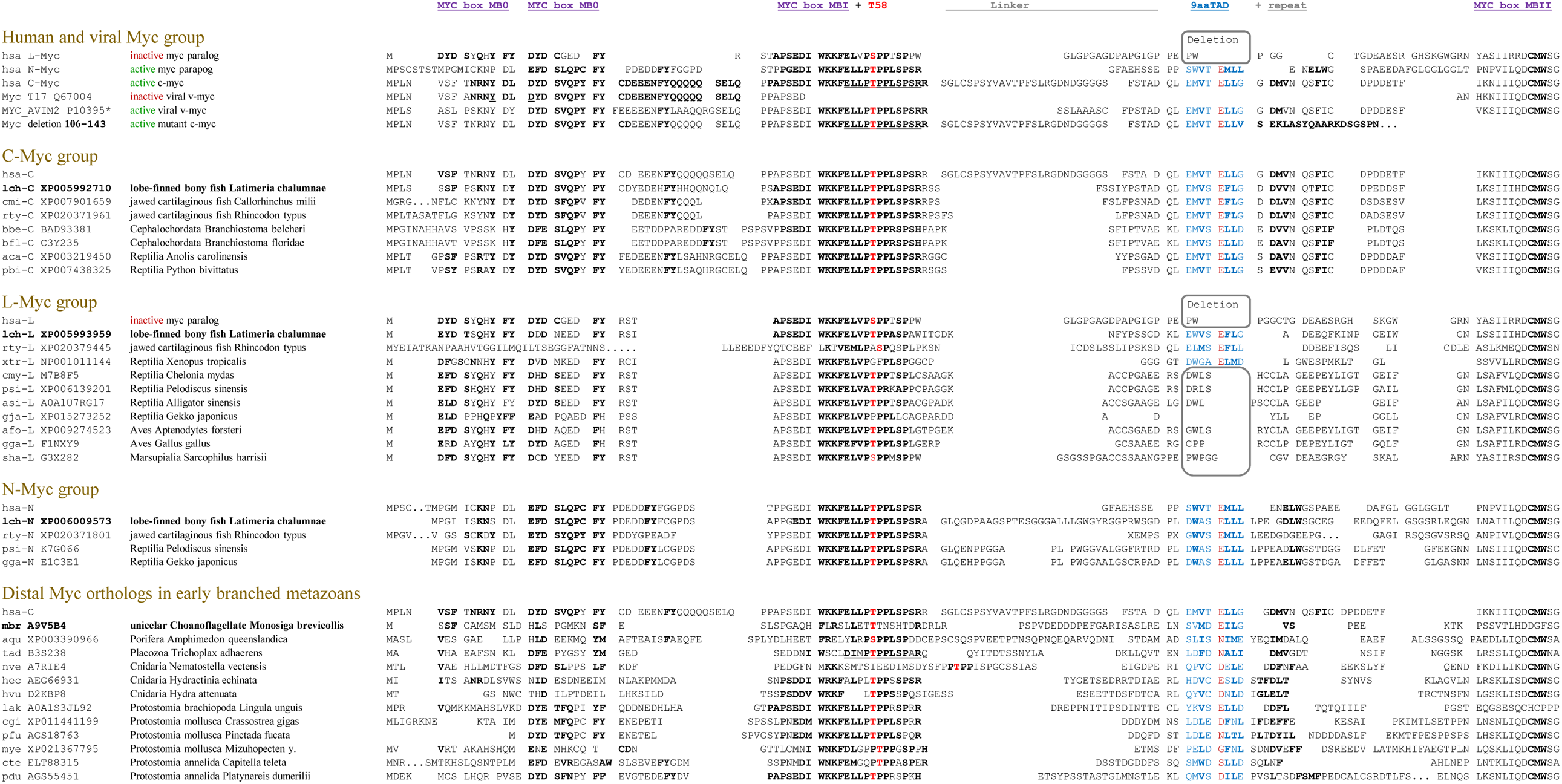
Alignment of the MYC family. The N-terminal regions of MYC proteins were aligned by sequence similarities and their predicted 9aaTAD activation domains are shown. The 9aaTADs activation domains are colored for faster orientation. A) Contrary to coelacanth (Latimeria chalumnae, fin-to-limb transition, related to lungfishes and tetrapods), the L-Myc from humans, reptiles and higher animals have no 9aaTAD motif and the corresponding region is deleted. B) The MYC clades with conservation MYB box O, I and II are shown. The deletion of activation domain in L clade are in grey box. Conservation of threonine in position 55 is highlighted in red. Pseudo repeat of the 9aaTAD motif with similarity to Sp2 9aaTAD is shown.

To identify the 9aaTAD activation domain, we ran our online 9aaTAD prediction service for trans-activating region 55-120 (localised between MYC boxes MBI and MBII) and revealed a 92% hit (sequence EMVT E LLGG, region 100-120) (**Figure 1**). Similarly, we revealed a 100% hit for the c-Myc mutant, which has an intact activation potential despite of the internal deletion (fusion result for modified trans-activation region with modified sequence EMVT E LLVS, fusion of regions 100-105 to 144-146).

We have analysed several previously reported mutants with surprising phenotypes. Reported insertion of glutamic acid into c-Myc (EMVT EL/ E /LVS, region 100-105 / inserted E / 144-146) (70) diminished the 9aaTAD pattern and its function. The activation domain 9aaTAD was also destroyed in mutants with deletion 3-103 and 106-143. Nevertheless, partial activity was observed in mutants with deleted regions 41-103, 56-103, 93-103 (including also small inserts due to DNA manipulation), which seem to have a partial repair of their 9aaTAD motif (QSEL / E LLGG, ELLD / E LLGG and GSSI / E LLGG).

The longest reported construct with full activity included the c-Myc region 1-262 (69). Despite of the presence of the activation region, a longer construct including region 1-336 has lost abruptly the capacity to activate transcription. Similarly, the mutant with deleted region 7-91 lost the capacity to activate transcription (70). Additionally, mutants with deleted regions 3-53 or 145-304 have no capacity to activate transcription regardless of their activation domains (regions 55-100). Together, we concluded that in these mutants with intact activation domains, the local structural aberrations hindered function of their activation domains.

The 9aaTAD activation domains are well conserved in the active human Myc paralog N-Myc, but completely absent in human paralog L-Myc, what well corresponds with its poor activation potential (71). Interestingly, we found conservation of the activation domains in all three Myc paralogs in lobe-finned bony fish coelacanth (*Latimeria chalumnae*, fin-to-limb transition, related to lungfishes and tetrapods) but not in reptiles and other higher metazoans, which suggests that loss of L-Myc activation capacity have occurred already in early tetrapod evolution (**Suppl. Figure S1**). Besides the L-Myc clade, we found conservation of the activation domains in the entire Myc family from human to the last unicellular ancestor of animals, the *Monosiga brevicollis* (72).

The regions coding for predicted activation domains were fused with the DNA binding domain of LexA in construct i) mbr Myc for flagellate filozoans *Monosiga brevicollis* (gene ID: A9V5B4), ii) construct aqu Myc for poriferans *Amphimedon queenslandica* (gene ID: XP003390966), iii) constructs lch c-Myc, lch N-Myc, lch L-Myc for lobe-finned fish *Latimeria chalumnae* (gene IDs: XP005992710, XP005993959, XP006009573), and iv) constructs hsa c-Myc and hsa N-Myc for humans. No construct was created for the human L-Myc, which has deleted activation domain and was reported as a poor activator (71). All tested Myc activation domains from human and coelacanth have the capacity to activate transcription (**Figure 2**). Previously, the accumulations of valines and isoleucines in activation domains of SP and KLF transcription factors were linked with their full or partial inhibition, which corresponds well to lower activities of mbr and aqu Myc constructs in our transactivation assay (22).

## Discussion

The activation domains have been reported as intrinsically disordered regions, which facilitate fuzzy binding with mediators of transcription. Their binding mechanism should be sequence independent because mutation of nearly every residue within is tolerated (23). In spite of enormous activation domain variability observed within the natural and also artificial activation domains, we generated a 9aaTAD pattern and prediction algorithm based on the amino acid distribution and clustering (18). In this and our other studies, the 9aaTAD online prediction (https://www.med.muni.cz/9aaTAD/index.php) generated reliable predictions, which led to the identification of over hundred activation domains, which have been experimentally confirmed as powerful activators of transcription. Moreover, the 9aaTAD structures seem to be helical after binding to mediators and their interaction with mediators were found not fussy but rather well fixed after binding e.g. MLL, E2A, Myb, p53 (1, 14, 73, 74). We also observed high similarity and partial identity in some of the 9aaTADs e.g. MLL and E2A (<u>SDI</u>M <u>D F</u>VLK - <u>SDL</u>L <u>D F</u>SAM) or human Myc and rat p53 (EM<u>V</u>T <u>E LL</u>G<u>G</u> - QD<u>V</u>A <u>E LL</u>E<u>G</u>) or human / Crassostrea gigas KLF4 and Oaf2 (<u>LLD</u>L <u>D FIL</u> / <u>LLDY D FIL</u> - <u>LFDY D FLF</u>)(14, 22). All that supports that 9aaTAD domains have a natural prevalence for some variants despite their tentatively huge variability.

In our study, we have identified activation domains in all four Yamanaka factors, Oct4, Sox2, Myc, and Klf4. All of them fulfil the criteria for the 9aaTAD domains, which activated transcription as short peptides. The loss of 9aaTAD in human L-Myc (as well as in reptiles and higher animals) explained its poor activator function and pointed to the functional diversification of MYC family during vertebrates’ evolution. The members of SOX clades B, F, C, E possess functional 9aaTADs, could partially substitute for each other and their function also largely diversified first in vertebrates. Similarly, the KLF family and their successor SP family (Klf1, Klf2, Klf3, Klf4, Klf5, Klf6, Klf7, Klf8, Klf12, Klf15, WT1, respectively Sp1, Sp2, Sp3, and Sp4), all possess a functional 9aaTAD and have versatile functions in cell fate determination.

We found the 9aaTAD pattern deterioration in Callorhinchus milii (Ghost shark) cmk Pit1 (Pou1 clade) indicating that the Pit1 9aaTAD activation domain is the latter evolution event in vertebrates. Similarly, Pou5 orthologs in Callorhinus ursinus (cur Pou5, Northern fur seal) and Sarcophilus harrisii (Tasmanian devil) have less transcriptional activity and even the loss of activity was found in Lingula unguis (brachiopod), which has also a deteriorated 9aaTAD motif. Moreover, we found a similar trend in Myc orthologs from flagellate filozoans Monosiga brevicollis and poriferans Amphimedon queenslandica, where the activation domains have substantially compromised function (disadvantageous accumulation of valines and isoleucines in their 9aaTADs, e.g. inactivated 9aaTAD in Sp2).

By cooperative interactions, active enhancers interact with mediators of transcription to enable assembly of the transcriptional machinery. The phase separation contributes to stabilization of transcription machinery at the specific genomic loci even at very low protein concentrations. Throughout the genome, the Oct4, Sox2 and Klf4 binding sites and mediators of transcription distribution have an exponential occurrence correlation, whereby the DNA binding sites serve as scaffolds for the phase-separated transcriptional condensate and cooperate during cell reprogramming. In summary, we consider the mediator interaction interface facilitated by the 9aaTADs in higher vertebrates as a part of Yamanaka factors ability to reprogram cell fate.

## Methods

### Constructs

The construct pBTM116-HA was generated by insertion of an HA cassette into the EcoRI site of the vector pBTM116. All constructs were generated by PCR and subcloned into the pBTM116 EcoRI and BamHI sites. All constructs had a spacer of three amino acids inserted into the EcoRI site: the peptide GSG. All constructs were sequenced by Eurofins Genomics. Further detailed information about constructs and primer sequences is available on request.

### Assessment of enzyme activities

β-galactosidase activity was determined in the yeast strain L40 [58, 59]. The strain L40 has a chromosomally integrated lacZ reporter driven by the lexA operator. In all hybrid assays, we used the 2μ vector pBTM116 for generation of LexA hybrids. The yeast strain L40, Saccharomyces cerevisiae genotype: MATa ade2 his3 leu2 trp1 LYS::lexA-HIS3 URA3::lexA-LacZ, is deposited at the ATCC (#MYA-3332). The average value of the β-galactosidase activities from three independent transformants is presented as a percentage of the reference with the standard deviations (means plus and minus SDs; n = 3). We standardized all results to the previously reported Gal4 construct HaY including the 9aaTAD with the activity set to 100% [14].

Databases used in the study. UniProt, ExPASy, NCBI, Ensembl Metazoa, KEGG http://www.genome.jp, Japanese Lamprey Genome Project http://jlampreygenome.imcb.a-star.edu.sg/blast/, Sun Yat-Sen University Lancelet Genome Project http://genome.bucm.edu.cn/lancelet/blast.php, Compagen Genomics Platform http://www.compagen.org/blast.html, Florida University Neurobase https://neurobase.rc.ufl.edu/, UCSC genomic annotation https://genome.ucsc.edu, NIH https://research.nhgri.nih.gov/mnemiopsis/blast/.

## Acknowledgements

This work was supported by the Ministry of Health of the Czech Republic AZV NV19-05-00410.

## Author contributions

MP, KJ, MR, and MH performed the experiments. MP conceived the project. MP and AK wrote the manuscript. All authors have contributed critical intellectual content and have approved the final manuscript.

